# Intratumoral MAPK and PI3K signaling pathway heterogeneity in glioblastoma tissue correlates with selective CREB signaling and specific functional gene signatures

**DOI:** 10.1101/305052

**Authors:** Paul M. Daniel, Gulay Filiz, Martin J. Tymms, Robert G. Ramsay, Andrew H. Kaye, Stanley S. Stylli, Theo Mantamadiotis

## Abstract

Limitations in discovering useful tumor biomarkers and drug targets is not only due to patient-to-patient differences but also due to intratumoral heterogeneity. Heterogeneity arises due to the genetic and epigenetic variation of tumor cells in response to microenvironmental interactions and cytotoxic therapy. We explored specific signaling pathway activation in glioblastoma (GBM) by investigating the intratumoral activation of the MAPK and PI3K pathways. We present data demonstrating a striking preponderance for mutual exclusivity of MAPK and PI3K activation in GBM tissue, where MAPK activation correlates with proliferation and transcription factor CREB activation and PI3K activation correlates with CD44 expression. Bioinformatic analysis of signaling and CREB-regulated target genes supports the immunohistochemical data, showing that the MAPK-CREB activation correlates with proliferative regions. In-silico analysis suggests that MAPK-CREB signaling activates a pro-inflammatory molecular signature and correlates with a mesenchymal GBM subtype profile, while PI3K-CREB activation correlates with the proneural GBM subtype and a tumor cell invasive gene signature. Overall, the data suggests the existence of intratumoral subtype heterogeneity in GBM and that using combinations of both MAPK and PI3K drug inhibitors is necessary for effective targeted therapy.

## Introduction

Problems associated with the treatment of recurrent glioblastoma (GBM) are thought to be due to variations in tumor cell drug sensitivity, as well as the development of tumor cell drug resistance, underpinned by the heterogeneous molecular and genetic nature of GBM. Unravelling the molecular mechanisms underlying the development of the most common and deadly malignant brain cancer, GBM, is necessary to define novel biomarkers and specific druggable targets. The biological and clinical differences of GBM are underpinned by the diverse underlying genetic and molecular landscape. The molecular heterogeneity in GBM is reflected by the varied phenotypic and functional features between GBM subtypes [29, 43]. There is significant difference in response to intensive treatment, which includes chemotherapy with temozolomide or concurrent chemotherapy and radiotherapy, among GBM subtypes. Proneural GBM is least responsive to intensive treatment while classical and mesenchymal subtypes respond relatively well [43]. Underlying differences in therapeutic response between each subtype may be due to the distinct regulatory networks driving the pathology, since each subtype has a unique regulatory network established by epigenomic, genomic and transcriptomic changes [4, 8]. As such, the divergent biological attributes of GBM subtypes make characterization not only important for prognosis but also relevant to identification of subtype-specific therapeutic targets.

Genomic analysis of the mutational spectra of GBM shows that mutations often occur in multiple components of key signaling pathways which lead to deregulation of downstream kinase cascades and drive tumorigenesis [6]. Hyperactivation of the MAPK pathway is common to many cancers including GBM and has been implicated in multiple tumorigenic processes such as proliferation [1], survival [18] and migration [14]. The core role of MAPK in promoting malignancy is highlighted by the correlation of hyperactivation of this pathway to poor prognosis in multiple cancer types including breast [27], colon [36], ovarian [32] and GBM [30, 31]. Signaling through this pathway begins at membrane bound growth factor receptors, typically the epidermal growth factor receptor (EGFR), which is activated by epidermal growth factor (EGF). Upon ligand binding, the cytoplasmic domain of the receptor becomes phosphorylated and recruits the downstream MAPK pathway signaling adapter proteins.

Clusters of mutations in multiple MAPK pathway components drive aberrant hyperactivation. In GBM, five of the top twenty-five mutated genes belong to the MAPK pathway and 83% of all GBM cases possess a mutation in at least one of these factors [13]. Of the mutations in the MAPK signaling pathway, the *EGFR* is one of the most common (50%) found in GBM. This is typified by the EGFRvIII activating mutation, a missense mutation which promotes ligand-independent activation of the receptor [19], resulting in the ligand-independent, constitutive activation of EGFR leading to hyperactivation of MAPK pathway. MAPK regulates various cellular processes via the activation of downstream transcription factors, which in turn regulate the expression of numerous target genes. A key transcription factors which MAPK regulates includes the cAMP Response Element Binding Protein (CREB) [46]. We previously showed that in GBM cells, CREB, a key regulator of cyclin-D1 expression cell and cell proliferation, is coactivated by MAPK and PI3K signaling [16].

Like the MAPK pathway, the PI3K pathway is aberrantly hyperactivated in multiple cancers, with 60-90% of GBM cases exhibiting an activating mutation in at least one of the core PI3K pathway genes [12, 20]. Activation of the PI3K pathway is associated with reduced survival in GBM patients [31] and regulates several characteristics of tumor cell biology including survival [11] and invasion [21]. Upon growth factor ligand-receptor interaction, the PI3K catalytic subunit, p110α, converts phosphatidylinositol 4,5-bisphosphate (PIP2) to phosphatidylinositol 3,4,5-triphosphate (PIP3), which then activates AKT [15]. AKT is the terminal kinase which has a host of target genes together with other kinases, including mTOR [23], pro-survival factor BCL2 [5] and transcription factors such as FOXO3 [38] and CREB [17]. Notably, many of the targets for the MAPK pathway are shared by the PI3K pathway, indicative of their overlapping roles in regulating tumor cell functions.

Mutations of activator or inhibitor proteins along the PI3K pathway create the net effect of PI3K pathway hyperactivation. For example, mutation of tumor suppressor and PI3K pathway negative regulator, phosphatase and tensin homologue (PTEN), is reported in 30-40% of GBM cases [12, 43]. Compared to the MAPK pathway, PI3K signaling activation correlates more weakly with prognosis in GBM [28, 31]. However, experimental evidence shows that PI3K pathway mutations are drivers / initiating events in glioma [41]. Several mouse models have also demonstrated that PTEN loss is necessary for initiating GBM when combined with other oncogenic mutations [2, 45].

The MAPK and PI3K cell signaling pathways were originally modelled as linear signaling intracellular conduits activated by distinct stimuli, but studies have since demonstrated that the pathways converge to regulate each other and co-regulate downstream functions (reviewed in [26]). However, each pathway also responds to and implements distinct molecular and cellular functions. With the view that targeting specific factors regulating specific pathways will disrupt key tumor cell behaviors, including survival and proliferation, we present data showing that in GBM tissue, there is regional variation with respect to MAPK and PI3K activation and that there is a striking preponderance for mutual exclusivity in the activation of the pathways. In situ coexpression analysis and computational analysis suggests that there are specific roles that each pathway plays in alliance with the CREB transcription factor. The intratumoral signaling pathway heterogeneity we identify, has implications on tumor cell behavior and likely has clinical implications on targeted therapy rationale and GBM subtype determination. Importantly, the data also supports the view that inhibiting both major cell signaling pathways is always necessary to effect tumor cell inhibition in GBM.

## Methods

### Ethics Statement

Experiments using fixed GBM tissue were carried out with the approval of The University of Melbourne Office for Research Ethics and Integrity, Human Research Ethics Committee, project ID 1339751.

### Immunostaining

Three different patient tissues, diagnosed as WHO Grade IV/GBM by pathologists at the Royal Melbourne Hospital, were used. Paraffin sections were cut at a thickness of 7μm, cleared of paraffin and rehydrated through an ethanol to water series. All staining and immunohistochemical analysis for each specimen was performed on consecutive serial sections. Hematoxylin and eosin staining was performed. Primary antibodies were incubated overnight at 4°C, washed in PBS and incubated with secondary antibody for 1h at ambient temperature, washed in PBS then developed for chromogenic or fluorescence visualization, as described in the following text. Immunohistochemical detection was performed using a Vectastain ABC kit (Vector Laboratories, Burlingame, CA) and diaminobenzidine (DAB; Merck, Darmstadt, Germany). Immunofluorescence staining was performed using secondary antibodies, Alexa-fluor 488 (green) or 594 (red), at 1:10000 dilution. Primary antibodies were: pAKT(S473) (#9271, Cell Signaling Technology, Danvers, MA), pMAPK pMAPK (#4370, Cell Signaling Technology, Danvers, MA), pCREB (#9198, Cell Signaling Technology, Danvers, MA), PCNA (#2686, Cell Signaling Technology, Danvers, MA), CD44 (#04-1123, Abcam, Australia), MMP-(#29579, Anaspec, Fremont, CA). Primary antibody dilutions were at 1:1000 for immunohistochemistry and 1:200 for immunofluorescence. For immunofluorescence staining, coverslips were mounted using Fluoroshield Mounting Medium with DAPI (Abcam, Australia).

### Pathway-specific CREB target gene analysis

The level 3 TCGA Agilent GBM dataset was utilized for the analysis (https://portal.gdc.cancer.gov/projects/TCGA-GBM). Gene sets (MSigDB) containing KEGG annotations for the MAPK and the PI3K pathways were used to perform single sample Gene Set Enrichment Analysis (ssGSEA) to define the level of pathway enrichment within individual patients in the GBM dataset. DEseq was used to test for differentially expressed genes between samples with high (z>1) and low (z<-1) enrichment of each KEGG pathway [3]. A false discovery rate corrected p-value of <0.1 was used as a cut-off for differentially expressed genes.

Pathway specific differentially expressed genes were then compared to each other to identify uniquely expressed genes specific to each pathway. To define CREB target genes, pathway specific genes were interrogated for the presence of a full CRE (cAMP response element) site in the promoter region, −1000bp to +100bp from the transcription start site (TSS), regions representing high confidence CREB target gene promoters [47]. The resulting genes were grouped as PI3K-CREB and MAPK-CREB gene modules for downstream analysis.

## Results

### Preponderance for intratumoral mutual exclusivity of MAPK and PI3K activation in GBM

Immunohistochemical analysis of consecutive formalin-fixed paraffin-embedded (FFPE) tissue sections from three independent GBM (WHO Grade IV) patient tissues showed that in regions enriched in pMAPK expression, pAKT expression was low/undetectable and vice versa (Fig 1A, B). pMAPK positive regions exhibited overlapping expression with proliferating cell nuclear antigen (PCNA), matrix metallopeptidase-9 (MMP-9) and pCREB, while regions high in pAKT expression overlapped with CD44 expression but not the other biomarkers. Olig-2 expression also closely correlated with pMAPK expression (not shown). Tissue immunofluorescence staining showed that the same cells expressing pMAPK also expressed pCREB (Fig 2A), confirming the immunohistochemical data, while pAKT positive cells were mostly pCREB negative (Fig 2B), although there was co-expression of pAKT and pCREB in the same cells in the bordering regions.

**Fig. 1.**
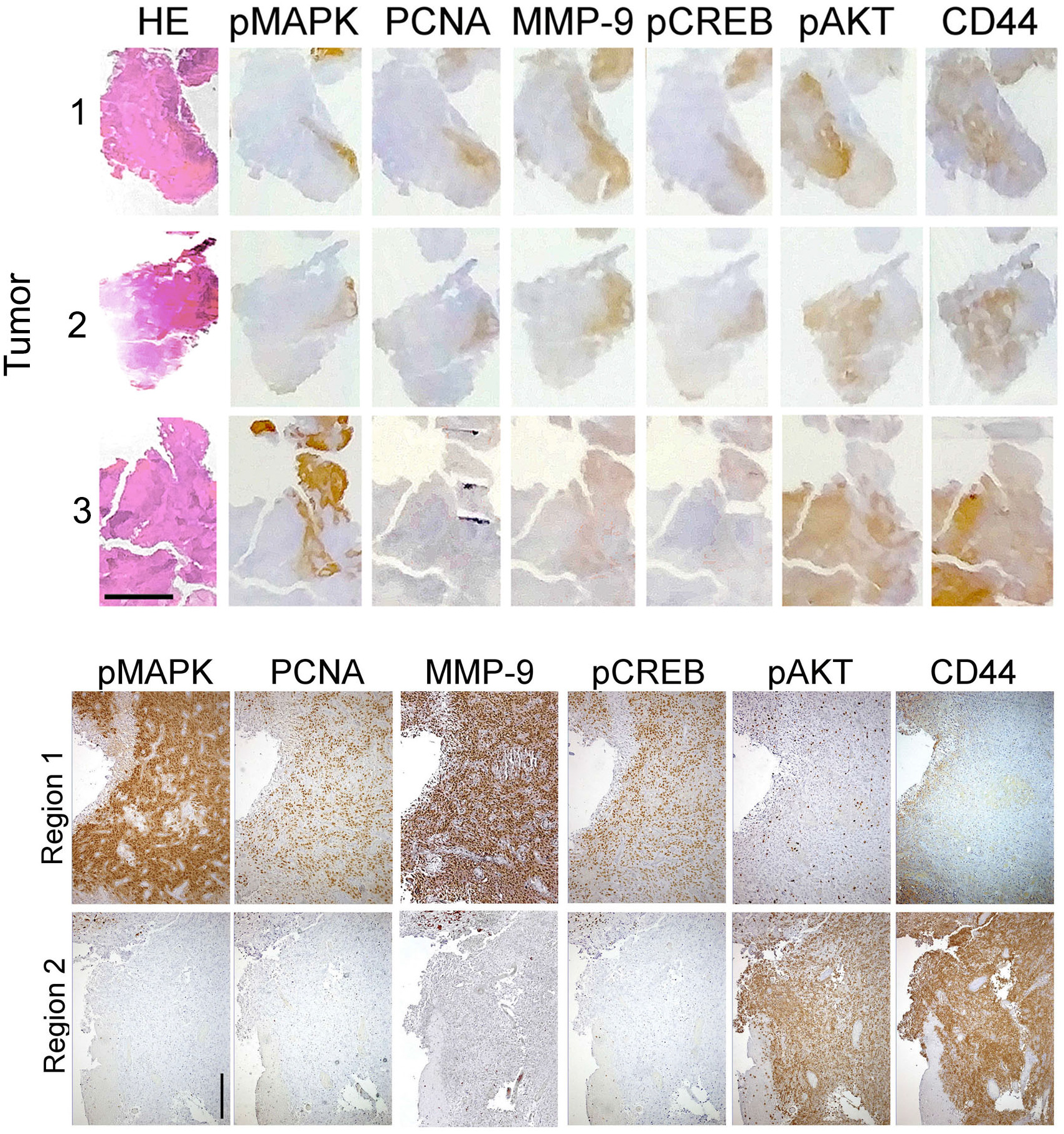
Distinct expression of activated MAPK and PI3K pathways and GBM tumor biomarkers in GBM tissue. (**A**) IHC analysis of groups of consecutive slides from three GBM patient specimens, showing distinct patterns of co-localization of pCREB with pERK1/2, pAKT(S473), PCNA and CD44. Scale bar = 5mm. (**B**) Biomarker expression in different regions of the same tumor (tumor 2). Scale bar = 200μm.

**Fig. 2.**
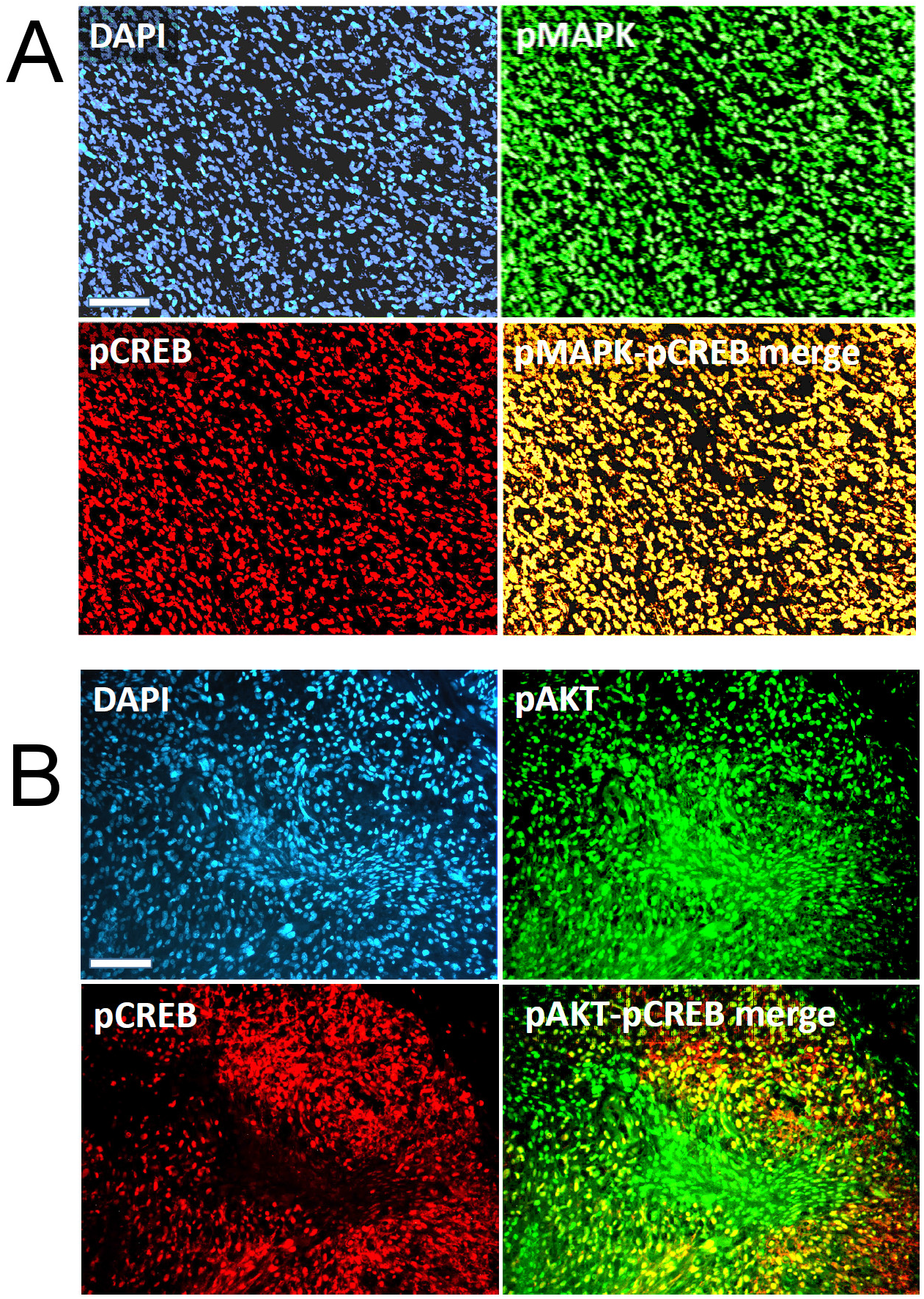
Co-expression of the MAPK and PI3K pathways with activated CREB in GBM tissue. Immunofluorescence of GBM patient specimens were co-stained with pAKT(S473) and pCREB antibodies or pMAPK and pCREB antibodies to investigate co-localization in GBM cells, showing the co-expression of these proteins in GBM tissue. Sections were also stained with DAPI to highlight the cell nuclei. Scale bar = 100μm.

### Distinct pathway-specific activation of CREB target gene expression in GBM

To investigate specific cell-associated functions of the MAPK and PI3K pathways in GBM and how these pathways differentially regulate common transcriptional targets, we used the TCGA GBM Agilent microarray RNA expression datasets and developed an in-silico analysis pipeline (Fig 3A), to identify CREB target genes expressed under the control of the PI3K pathway or MAPK pathway. The expression of 132 CREB target genes were specifically expressed when the MAPK pathway was activated (MAPK-CREB gene module) while 114 CREB target genes were expressed, only when the PI3K pathway was activated (PI3K-CREB gene module) (Table 1).

**Fig. 3.**
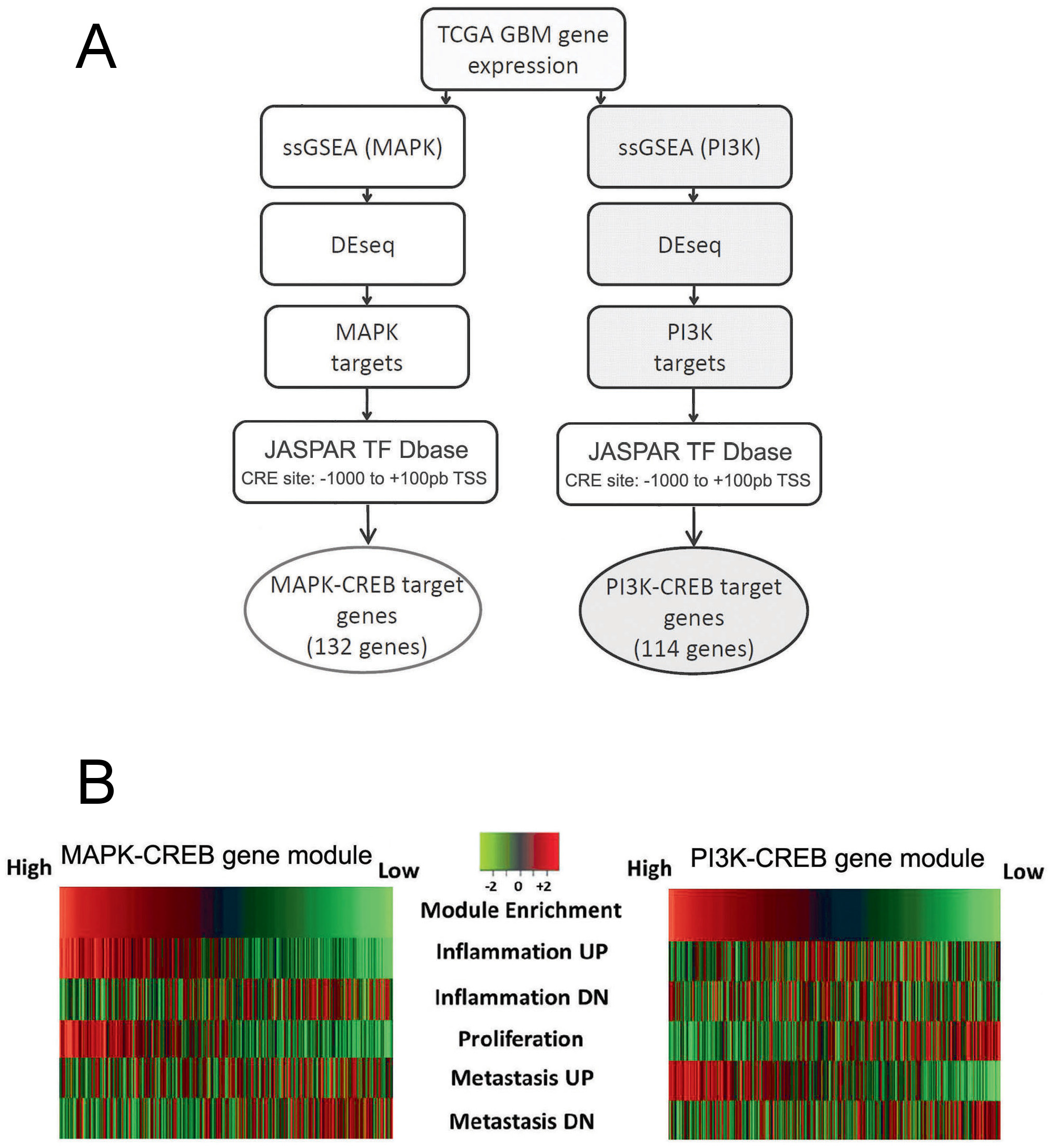
Pathway-specific CREB target genes are functionally distinct in GBM. A) Schematic showing the flow of single-sample Gene Set Enrichment Analysis (ssGSEA) used to analyze the TCGA GBM gene expression data set to enrich for MAPK and PI3K signature genes which are putative CREB target genes. See Methods for details. (B) GSEA analysis of 543 GBM patient expression and clinical data for signal pathway and CREB target gene enrichment demonstrating enrichment of MAPK-CREB genes and PI3K-CREB genes and associated tumor functions; colored bars represent individual patients from the TCGA cohort.

**Table 1.** CREB targets regulated by the MAPK and PI3K pathways. The number of genes differentially expressed in GBM (TCGA-GBM dataset), compared to normal brain tissue. Genes were divided into those associated with either the MAPK or PI3K pathways, using the Kyoto Encyclopedia of Genes and Genomes (KEGG) gene function annotations. From these genes, overexpressed genes were selected and CREB target genes identified. The last column indicates the number of CREB target genes unique to each pathway.

| Pathway (KEGG) | Differentially Expressed Genes | Overexpressed Genes | Total CREB target genes | CREB Target Genes Unique to Pathway |
| --- | --- | --- | --- | --- |
| MAPK Pathway | 2846 | 930 | 161 | 132 |
| PI3K Pathway | 2546 | 1068 | 203 | 114 |

**Table 2.** Gene ontology for PI3K-CREB and MAPK-CREB target gene modules. Listing of the most significant genes identified using the pipeline analysis described in Material and Methods and Fig 3A. (FDR = False-Discovery Rate).

| Gene module | Gene Ontology<br>Descriptor | Gene Symbol | p-value | FDR |
| --- | --- | --- | --- | --- |
| PI3K-CREB | Cellular<br>Macromolecule<br>Metabolic Process | PICK1, PDPK1, CDC42BPB,<br>GAL3ST1, DUSP8, CSNK1E,<br>TUFM, B3GALT2, UBE4A2,<br>PYGM, MAPT, EIF4A2,<br>MGEA5 | $4.49 \times 10^{-6}$ | $6.53 \times 10^{-3}$ |
| | Biopolymer<br>Metabolic Process | PICK1, PDPK1, CDC42BPB,<br>GAL3ST1, DUSP8, CSNK1E,<br>TUFM, B3GALT2, UBE4A2,<br>ARID1A, HDAC5, SNR,<br>HTATSF1, KLF13 | $1.69 \times 10^{-5}$ | $6.59 \times 10^{-3}$ |
| | Cellular Protein<br>Metabolic Process | PICK1, PDPK1, CDC42BPB,<br>GAL3ST1, DUSP8, CSNK1E,<br>TUFM, B3GALT2, UBE4A2,<br>MAPT, EIF4A2, MGEA5 | $2.08 \times 10^{-5}$ | $6.59 \times 10^{-3}$ |
| MAPK-CREB | Positive Regulation<br>of Biological Process | SCG2, NLRC4, TNFSF14, PML,<br>ECL1, DAPK3, TICAM1,<br>RHOG, VEGFA, MAP2K3,<br>HSP90AA1, TPP1, FLT1, C2 | $1.98 \times 10^{-8}$ | $2.85 \times 10^{-5}$ |
| | Signal Transduction | SCG2, NLRC4, TNFSF14, PML,<br>ECL1, DAPK3, TICAM1,<br>RHOG, VEGFA, MAP2K3,<br>ARHGAP27, GADD45B,<br>TNFAIP3, FGD5, HOMER3,<br>TRH, ZYX, CSNK1G2, IL7R,<br>RCAN1 | $4.86 \times 10^{-8}$ | $3.54 \times 10^{-5}$ |
| | Positive Regulation<br>of Cellular Process | SCG2, NLRC4, TNFSF14, PML,<br>ECL1, DAPK3, TICAM1,<br>RHOG, VEGFA, MAP2K3,<br>HSP90AA1, TPP1, FLT1, | $7.64 \times 10^{-8}$ | $3.7 \times 10^{-5}$ |

Gene clustering analysis showed that PI3K-CREB signaling regulates target genes identified by the GO (Gene Ontology) term ‘metastasis’, suggesting that these genes are involved in regulating tumor cell migration / invasion, while MAPK-CREB regulated genes are categorized as genes regulating ‘proliferation’ and ‘inflammation’ (Fig 3B). Further interrogation of genes showed that MAPK-CREB genes were enriched for GO terms, ‘Positive Regulation of Biological Process’, ‘Signal Transduction’ and ‘Positive Regulation of Cellular Process’, whereas PI3K-CREB genes were enriched for GO terms ‘Cellular Macromolecule Metabolic Process’, ‘Biopolymer Metabolic Process’ and ‘Cellular Protein Metabolic Process’. Further curation of gene function using Genecards (http://www.genecards.org/) [34], identified specific cellular and molecular functions of the target genes and identified genes which may be involved in cross-pathway regulation (Table 3). PI3K-CREB regulated, DUSP8, negatively regulates MAPKs, while the MAPK-CREB regulated protein PML inhibits AKT1, a key kinase of the PI3K pathway[34]. Overall, PI3K-CREB regulated genes were themselves regulators of multiple signal transduction pathways, including the PI3K, WNT, JNK, PKC and MAPK pathways and regulators of transcription, illustrating the broad involvement of the PI3K-CREB in regulating complex molecular and cellular functions. The MAPK-CREB target genes identified also fell into broad categories but also identified inflammation and angiogenesis as key, more specific functions (Table 3).

**Table 3.** Molecular functions for PI3K-CREB and MAPK-CREB target genes. Functions were identified by GO in Table 2. Bold gene names indicate factors with cross-pathway regulatory functions.

| Gene module | Gene Symbol | Molecular Function | Cell Function |
| --- | --- | --- | --- |
| PI3K-CREB | PDPK1 | PI3K PATHWAY | Signaling |
|  | CSNK1E | WNT PATHWAY |  |
|  | CDC42BPB | JNK PATHWAY |  |
|  | PICK1 | PKC PATHWAY |  |
|  | **DUSP8** | **MAPK SIGNALING** |  |
|  | ARID1A<br>HDAC5 | CHROMATIN REMODELLING | Transcription |
|  | HTATSF1 | ELONGATION FACTOR |  |
|  | KLF13 | REPRESSOR |  |
| MAPK-CREB | NLRC4 | INFLAMMASOME | Inflammation |
|  | ECL1 | CHEMOKINE RECEPTOR |  |
|  | TNFAIP3 | TNF PATHWAY |  |
|  | TNFSF14<br>TNFAIP3<br>TICAM1 | NFKB PATHWAY | Signaling |
|  | MAP2K3 | MAPK PATHWAY |  |
|  | **PML** | **PI3K INHIBITOR** |  |
|  | CSNK1G2 | WNT PATHWAY |  |
|  | HSP90AA1 | CHROMATIN MODELLING | Transcription |
|  | VEGFA | GROWTH FACTOR | Angiogenesis |
|  | FLT1 | VEGF-A/-B RECEPTOR |  |
|  | FGD5 | RAS-RELATED PROTEIN |  |
|  | VEGFA | GROWTH FACTOR | Cell migration |
|  | FLT1 | VEGF-A/-B RECEPTOR |  |
|  | FGD5 | RAS-RELATED PROTEIN |  |
|  | RHOG | RAS HOMOLOG |  |
|  | DAPK3 | CASPASE PATHWAY | Apoptosis |
|  | TICAM1 | TOLL-LIKE RECEPTOR |  |
|  | GADD45B | DNA DAMAGE RESPONSE |  |

### MAPK-CREB signaling correlates with proliferation and inflammation

For patient tumors highly enriched for MAPK-CREB gene expression, there was a strong correlation with proliferation and pro-inflammatory functions (Fig 3). The 115 gene PI3K-CREB module, showed an enrichment of genes involved in metastasis but no association with proliferation or pro-inflammation genes. VEGFA, FLT1 and FGD5 were amongst the genes identified which regulate proliferation and angiogenesis, while three genes involved in the NFκB pro-inflammatory pathway, TNFS14, TNFAIP3 and TICAM1, were also identified as MAPK-CREB regulated genes (Table 3).

### MAPK-CREB target genes but not PI3K-CREB target genes are associated with poorer survival in GBM

To determine whether the difference in gene targets regulated by the MAPK-CREB and PI3K-CREB modules was reflected in patient prognosis, clinical parameters from the TCGA GBM dataset were analyzed (Fig 3-5A, B). Patient tumors which were enriched for MAPK-CREB target genes had a poorer prognosis than those which were not enriched for this gene module (372 days vs 502 days, p=6.21×10^−4^; Fig 4A). By contrast, PI3K-CREB target gene expression did not correlate with a difference in survival between patients which had high or low gene enrichment scores. Notably, PI3K or MAPK gene expression modules, independent of the CREB-regulated target genes with high-confidence CRE promoter sequences, did not identify populations with significantly different survival (not shown).

**Fig. 4.**
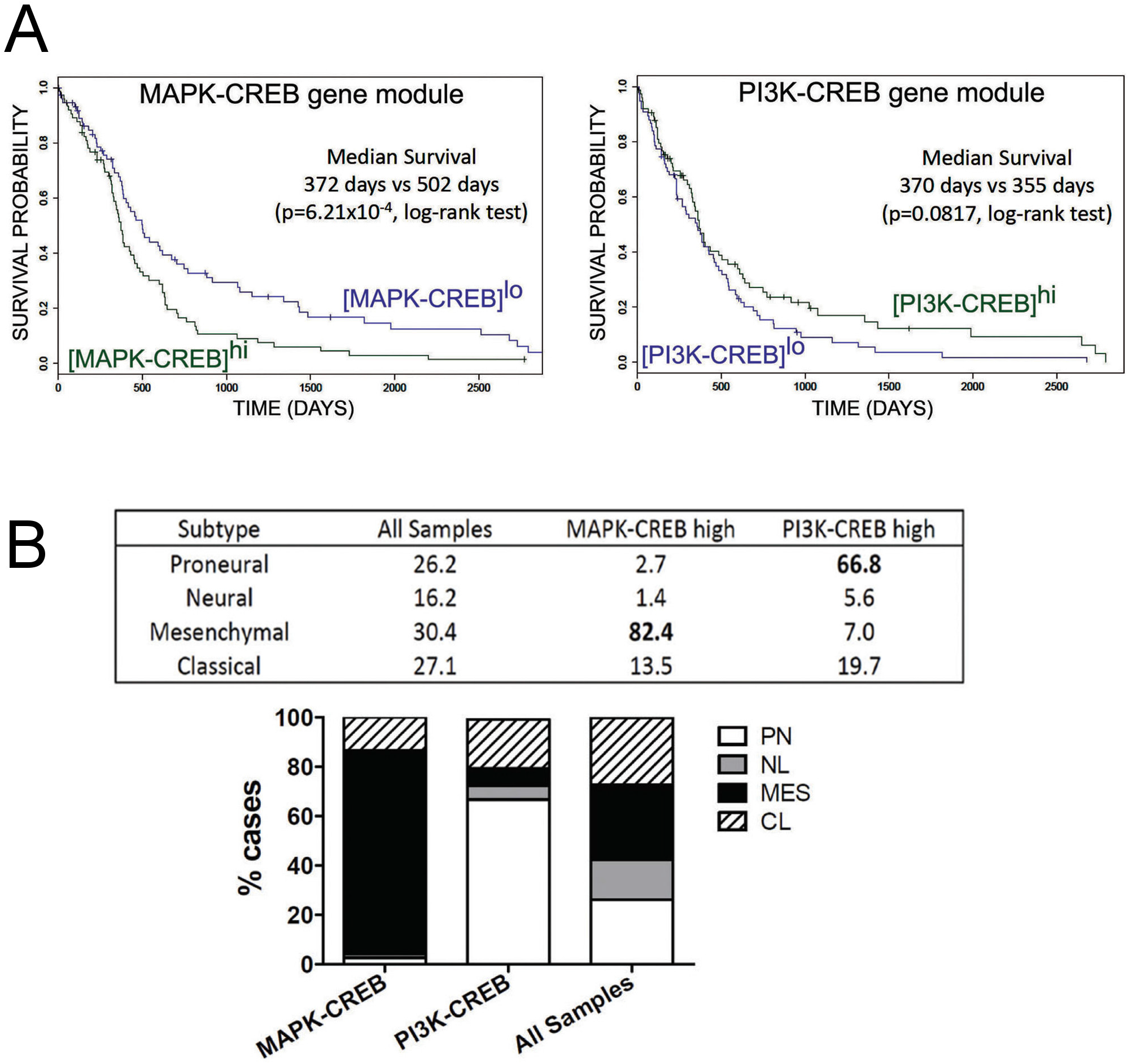
Pathway-specific CREB target gene expression correlates with survival and GBM subtype. A) Survival analysis of patients defined by enrichment of MAPK-CREB and PI3K-CREB target genes. B) The MAPK-CREB signaling-transcription module associates with Mesenchymal (MES) subtype GBM and PI3K-CREB module associates with Proneural (PN) GBM. “All samples” refers to the GBM subtype distribution of the 543 patient samples in TCGA. NL – Neural; CL – Classical.

### Pathway-specific CREB transcriptomes correlates with GBM subtype

Previous data, from deep genomic and proteomic analysis of 543 GBMs, has implicated distinct signaling pathways to be represented in each GBM subtype; with MEK/MAPK highly activated the mesenchymal subtype and PI3K highly activated in the proneural subtype [7]. To investigate whether the respective PI3K-CREB and MAPK-CREB gene expression modules represented distinct GBM subtypes, computational analysis was undertaken using the TCGA GBM dataset which includes information on the GBM subtype per patient specimen. The mesenchymal subtype expression profile was significantly over-represented (p=6.7−10^−8^) in GBM enriched for the MAPK-CREB module, while the PI3K-CREB module was enriched in the proneural subtype (Fig 4B), consistent with earlier work [7].

## Discussion

The MAPK and PI3K signal transduction pathways are aberrantly activated in cancer cells and this aberration drives GBM drives key tumor cell behaviors advantageous to these cells, including proliferation, survival and migration/invasion. The mutual exclusivity between the PI3K and MAPK pathways identified in our study is poorly described in the literature. In vitro investigation of signal transduction often reports that the MAPK and PI3K pathways are coactivated and are somewhat redundant and are also able to compensate for each other [44]. One contributing factor to this discrepancy between in vitro and in vivo results we present, is the intrinsic bias which culture conditions have on signaling. Cell lines are usually grown as a monolayer and maintained in medium containing serum which ensures every cell is exposed to the same growth factors which simultaneously hyperactivate multiple pathways to maximize cell proliferation and survival. This does not replicate the diverse and dynamic conditions of the tumor microenvironment, with heterogeneous tumor cell activation, which over time, regulates distinct tumor phenotypes.

Tumor heterogeneity is largely attributed to genomic events including mutations and epigenetic changes within tumor cells. Moreover, many targeted therapies attempt to inhibit one of these two key pathways using drugs targeting one of more pivotal molecules within the pathway. One of the difficulties in the analysis of tumor biology is a whole-tumor molecular view of intracellular events. Various experimental techniques can determine tumor gene sequence, mRNA and/or protein expression levels [12, 25, 43]. Examination of large numbers of patient GBM tissue biopsies using tissue microarrays have yielded important data on the expression of hundreds of proteins [42]. Despite the importance of these large-scale “omic” studies, the samples are lysates from a mixed cell population, devoid of the regional heterogeneity in tumors, in situ. High-throughput single cell analysis techniques can bypass some of the issues, but it remains to be seen how representative individual isolated cell experiments are, compared to in situ cell behaviors. One approach to interrogate the molecular events in GBM is to examine large tissue sections and multiple biomarkers which represent specific biological functions and/or therapeutic targets. A recent study demonstrates the value of this approach and analyzed tumor biomarkers using an integrated multiplexed tissue imaging platform on FFPE breast tumor tissue and correlated biomarker expression and MAPK and PI3K pathway activation with specific oncogenic mutations and histopathological diagnosis and demonstrated regional intra-tumoral pathway and biomarker expression heterogeneity [35]. We previously showed that proliferative and invasive GBM tumor regions, characterized by high olig-2 and CD44 expression, respectively, are regionally separated [9, 10] and that this expression is also associated with distinct GBM subtypes and that GBM heterogeneity can at least, in part, be attributed to intrinsic properties of the tumor cells. However, the status of the MAPK and PI3K pathways was not investigated.

In the present study, we showed that in GBM there was a mutual exclusivity of MAPK active tissue regions and PI3K active regions, in the same tissue biopsy. The activation status of MAPK active regions correlated with expression of the proliferation biomarker, PCNA [39], tumor cell invasion biomarker MMP-9 [22], pCREB expression, a pro-oncogenic transcription factor regulating genes involved in proliferation and survival [16, 24, 37] but little or no expression of CD44, a biomarker of tumor stem cells and cell invasion [33]. PI3K pathway active regions exhibited high levels of CD44 expression but little or no expression of PCNA, pCREB or MMP-9. The regionally distinct expression of MMP-9 and CD44 is interesting, since both genes are associated with tumor cell invasion/migration but in this case, it appears that these proteins may have distinct tumor promoting functions.

In-silico analysis showed that CREB sits at signaling convergence point downstream of both the MAPK and PI3K pathways with target genes activated common to both pathways but also distinct target genes, specific for each of the upstream pathways. Highly expressed PI3K-CREB target genes are prominent in ‘metastasis’ functions, suggesting that PI3K-CREB signals may be important for regulating the invasive behaviors of GBM cells. MAPK-CREB target genes are clearly associated with proliferation and inflammation. The clinical relevance of the analysis was highlighted by the finding that MAPK-CREB target genes but not PI3K-CREB target genes are associated with poorer survival in GBM and that the mesenchymal GBM subtype was overrepresented in patient tissue showing high MAPK-CREB target genes expression, whereas PI3K-CREB target gene expression correlated with the proneural GBM subtype. It is important to note that the distinct mesenchymal and proneural subtypes discussed in our study coexist in the same tumor tissue, while the literature has generally inferred that GBM subtypes are patient-specific. Our findings support a less prevalent view that single GBM tumors harbor more than one subtype signature, as proposed by Sotoriva and colleagues [40]. This also raises the question of, why the distinct signaling-CREB gene modules show differences in patient survival? We propose that when MAPK-CREB target gene expression across the whole GBM tumor is high, this drives a more malignant phenotype and shorter survival, compared with tumors with higher PI3K-CREB regulated gene expression.

Overall, the data presented here suggests that distinct MAPK and PI3K pathway activation and the downstream consequences lead to specific tumor cell responses. These regional variations may be caused by transient or long-term microenvironmental differences across the tissue. Regional signaling responses in the tumor tissue may also account for setting up topographically defined tumor tissue domains, similar to morphogenetic field responses in embryonic tissue. It is possible that neighboring cells within GBM tissue exhibit transient dynamic signaling events but how dynamic these regional differences are, remains to be determined and would require a means to monitor pathway activity, in situ. Another important issue raised by the data, is that GBM tumor cells do not require the simultaneous activation of both the MAPK and PI3K pathways. The clinical significance of the study is that it supports the view that for effective targeted GBM therapy, inhibiting both pathways, in combination with standard therapy, is necessary. If combined targeting proves to be too toxic, an alternative strategy may be that pharmacologically manipulating tumor cell signaling and dependence, from MAPK to PI3K, may shift GBM cells to a less malignant state, extending he therapeutic window and overall survival.

## Acknowledgments

Parts of this work were funded by a CASS Foundation Grant (7941) and Brain Foundation Award.

